# Climate and ant richness explain the global distribution of ant-plant mutualisms

**DOI:** 10.1101/2022.05.07.490958

**Authors:** Yangqing Luo, Amanda Taylor, Patrick Weigelt, Benoit Guénard, Evan P. Economo, Arkadiusz Nowak, Inderjit, Holger Kreft

## Abstract

Biotic interactions are known to play an important role in shaping species geographic distributions and diversity patterns. However, the role of mutualistic interactions in shaping global diversity patterns remains poorly quantified, particularly with respect to interactions with invertebrates. Moreover, it is unclear how the nature of different mutualisms interacts with abiotic drivers and affects diversity patterns of mutualistic organisms. Here, we present a global-scale biogeographic analysis of three different ant-plant mutualisms, differentiating between plants bearing domatia, extrafloral nectaries (EFNs), and elaiosomes, based on comprehensive geographic distributions of ∼15,000 flowering plants and ∼13,000 ant species. Domatia and extrafloral nectaries involve indirect plant defenses provided by ants, while elaiosomes attract ants to disperse seeds. Our results show distinct biogeographic patterns of different ant-plant mutualisms, with domatium- and EFN-bearing plant richness decreasing sharply from the equator towards the poles, while elaiosome-bearing plants prevail at mid-latitudes. Contemporary climate, especially mean annual temperature and precipitation, emerge as the most important predictor of ant-associated plant diversity. In hot and moist regions, typically the tropics, domatium- and EFN-bearing plant richness increases with related ant guild richness, while in warm regions plants with elaiosomes are strongly linked to interacting ants. Our results suggest that ant richness in combination with climate drives the spatial variation of plants bearing domatia, extrafloral nectaries, and elaiosomes, highlighting the importance of mutualistic interactions for understanding plant biogeography and its response to global change.

## Introduction

Mutualisms are fundamental for understanding the origin and maintenance of biodiversity (1, 2). Interacting species have evolved various traits that serve key functions in mutualistic interactions (2). Geographic variation of interacting species and related traits can provide insights into the drivers governing spatial patterns of biodiversity and species responses to future change (3, 4). It is especially true when considering the influences of biotic interactions on the distribution of interacting species at local spatial scales (5). However, a growing body of research suggests that biotic drivers can influence broad-scale patterns of mutualistic species and related traits, such as seed dispersal mutualisms (6, 7) and plant-mycorrhizal fungi mutualisms (8). Despite increasing evidence on spatial variation in diversity explained by biotic interactions, the relative importance of biotic and abiotic drivers on species diversity for plant-invertebrate mutualisms remains largely unquantified at macroecological scales, which are critical for understanding global plant diversity patterns.

Flowering plants and ants have evolved diverse mutualistic interactions, which arose from the evolution of plant adaptive traits to ant exploitation and eventually enhanced diversification in related plant lineages (2, 9–12). Over 15,000 plant species rely on ants as defenders and seed dispersal vectors, with the most common plant structures being domatia, extrafloral nectaries (EFNs) and elaiosomes (11–13). Domatia are modified plant structures such as cavities in leaves, stems, or roots that house ant colonies (13). Extrafloral nectaries are plant glands that attract ants by secreting sugary nectar outside of flowers (12). Ants, in return, protect domatium- and EFN-bearing plants against herbivores (14). On the other hand, elaiosomes are nutrient-rich appendages of seeds, which reward ant mutualists that disperse seeds (myrmecochory) (11). Plants interacting with ants have been studied in a variety of habitats and vegetation types, and at local to regional scales (10, 15). Consequently, ants are expected to shape the composition and structure of plant communities. For instance, plants with EFNs tend to be scarce on isolated islands that lack native ants compared to mainland floras (16). The “ant limitation hypothesis” postulates that ant community attributes, including ant species richness and forager abundance, may contribute to the evolutionary origin and maintenance of ant defense mechanisms in plants (16, 17). Likewise, ant activity and species richness are considered to affect myrmecochorous plant species diversity (18, 19). Accordingly, ant diversity may act as a biotic driver in shaping plant diversity patterns, similar to previously observed cross–taxon congruences in diversity pattern (20, 21).

Despite close associations of ecological and evolutionary histories between ants and plants, the diversity relationship between them is not always evident (22), suggesting alternative mechanisms may be more important. Abiotic factors, such as climate, may be responsible for geographic patterns of ant-associated plants. For example, the maintenance of ant defenses and dispersal could be influenced by resource availability. Productive habitats and regions may promote ant defenses, where the energetic costs of producing domatia and secreting sugary nectar are relatively inexpensive while the protection of fresh young leaves is essential (“resource limitation hypothesis”) (17, 23). The efficiency of ant defenses also depends on daily fluctuations in temperature, as ants behave more aggressively in warm periods (24). In contrast to ant-defended plants, plants with elaiosomes attached to their seeds typically occur in dry and low-nutrient soils (e.g., Mediterranean climate in Australia and South Africa). Indeed, myrmecochory is more affordable and directional in contrast to vertebrate dispersal, as seeds are protected in humid and nutrient-rich ant nests instead of above ground (2, 9).

Reflecting its fundamental importance for biodiversity, a compelling body of research has accumulated on the drivers of ant-associated plant diversity at local scales. However, our understanding of the relative importance of abiotic and biotic factors at larger scales is very limited, and historically hampered by data deficiency of the interacting species, particularly for invertebrate groups such as ants. Here, by integrating global distribution datasets of plants and ants, as well as checklists of plants with ant-related, specialized defense and dispersal traits (13, 25–28), we present global patterns of the three main ant-plant mutualisms, namely domatium-, EFN- and elaiosome-bearing plants. Specifically, we assess whether species richness patterns of ant-associated plants are driven by interacting ant richness, contemporary climate, and paleoclimate. Our study provides novel insights into the effects of biotic and abiotic factors on biodiversity in relation to three different ant-plant mutualisms.

## Results

### Geography of ant-associated plants

The three different groups of ant-associated plants show markedly distinct geographic patterns in species richness and proportional representation (relative proportion of interacting plants to all angiosperms per region) (Fig. 1 and *SI Appendix*, Fig. S1). Plants with domatia and EFNs exhibit marked latitudinal gradients with species richness and proportion peaking in tropical regions (Fig. 1). The highest diversity (domatia: 118 species in Peru; EFNs: 513 species in Bolivia) and proportion (domatia: 2.5%; EFNs: 19% of all angiosperms in Colombian departments, e.g., Guainía and Sucre) are typically found in tropical rainforests (*SI Appendix*, Table S1). Compared to widely distributed EFNs that show moderate species richness in temperate regions, domatia are almost exclusively found in tropical regions. Elaiosome-bearing plant richness, in contrast, peaks in the subtropics, decreasing towards both the poles and the tropics (Fig. 1). Hot, semi-arid regions contain the highest species richness (1528 species in South Africa) and proportion (17% species in South Western Australia) of plants with elaiosomes (*SI Appendix*, Fig. S2 and Table S1). Mediterranean regions (e.g., 511 and 9.1% species in Spain mainland, 9% species in Greece) and temperate forests of the northern hemisphere (e.g., 684 species in Sichuan, China) are also rich in elaiosome-bearing plants (*SI Appendix*, Tables S1).

**Figure 1.**
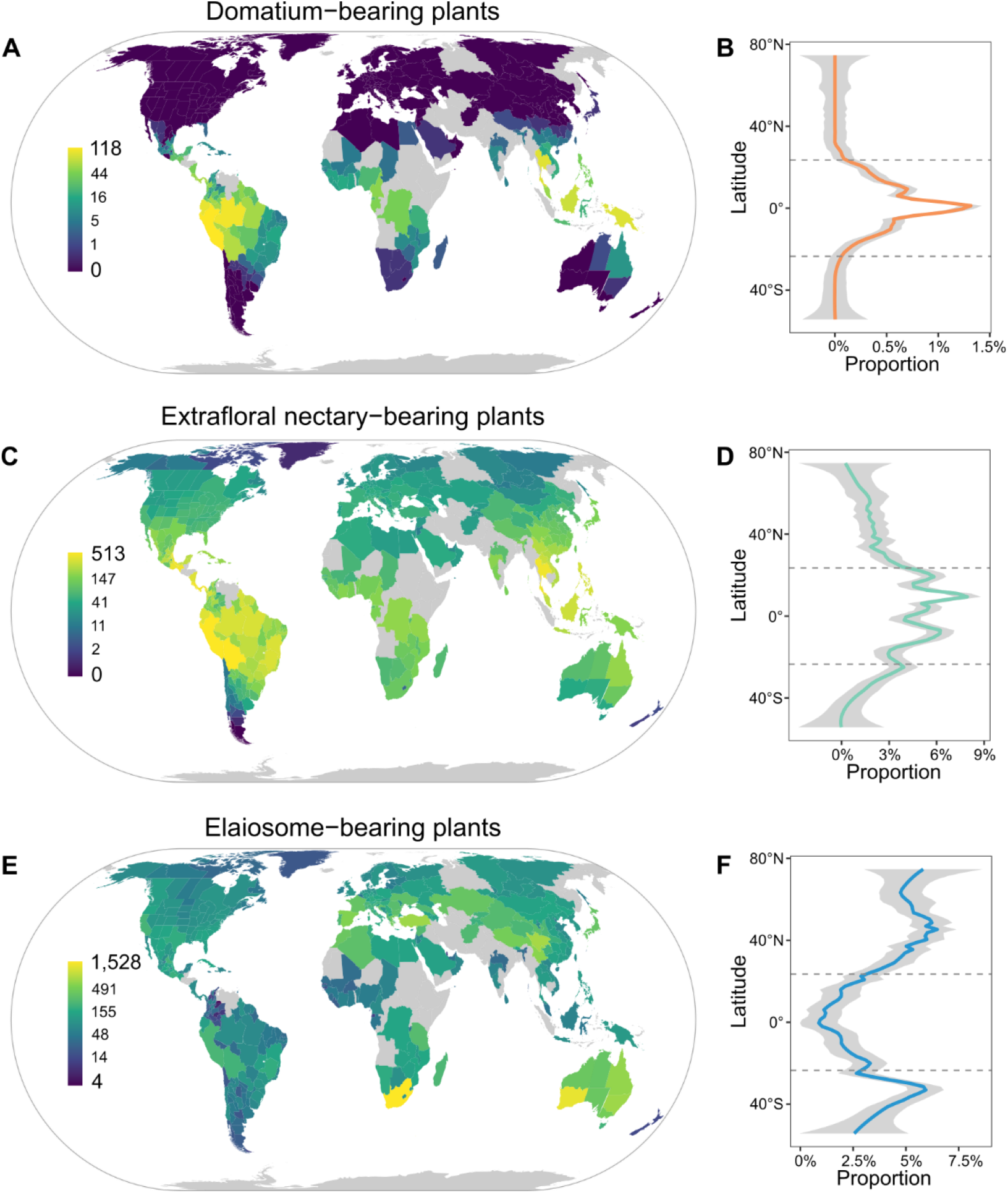
Global patterns of species richness and proportional representation for (*A* and *B*) domatium-bearing plants, (*C* and *D*) extrafloral nectary-bearing plants, (*E* and *F*) elaiosome-bearing plants. Species richness is indicated by the color of regions. Gray color indicates regions without data. Proportional representation is estimated as species richness of ant-associated plants relative to angiosperms and indicated along the latitudinal gradient.

### Biotic and abiotic drivers

To assess the direct and indirect biogeographic drivers of ant-associated plants, we performed structural equation modeling (SEM), including climatic (mean annual temperature, mean annual precipitation, precipitation seasonality, past climate change velocity in temperature and precipitation from the Last Glacial Maximum ∼21,000 y ago) and geographic variables (area, elevation range) as abiotic factors, and species richness of respective ant guilds as biotic factors. Ant guilds were classified as ants that may interact with domatia, EFNs and elaiosomes, respectively, based on an exhaustive literature search and expert opinions (*SI Datasets* S1). We found that species richness of ant-associated plants at broad spatial scales is strongly influenced by species richness of respective ant guilds, climatic, and geographic variables (Fig. 2). Temperature is positively related to species richness of plants with domatia and EFNs, while negatively related to plants with elaiosomes. On the other hand, precipitation exhibits a positive relationship with all three plant groups, but the effect on elaiosome-bearing plants is weak (Fig. 2). Compared to contemporary climatic variables, past climate change has only marginal effects (Figs. 2 and 3). Regions with higher climate change velocity in temperature are associated with lower species richness of domatium-bearing plants, and velocity in precipitation is negatively associated with EFN-bearing plants. Additionally, area shows strong direct effects on species richness of each plant group, while elevation range has only a low importance (Figs. 2 and 3).

**Figure 2.**
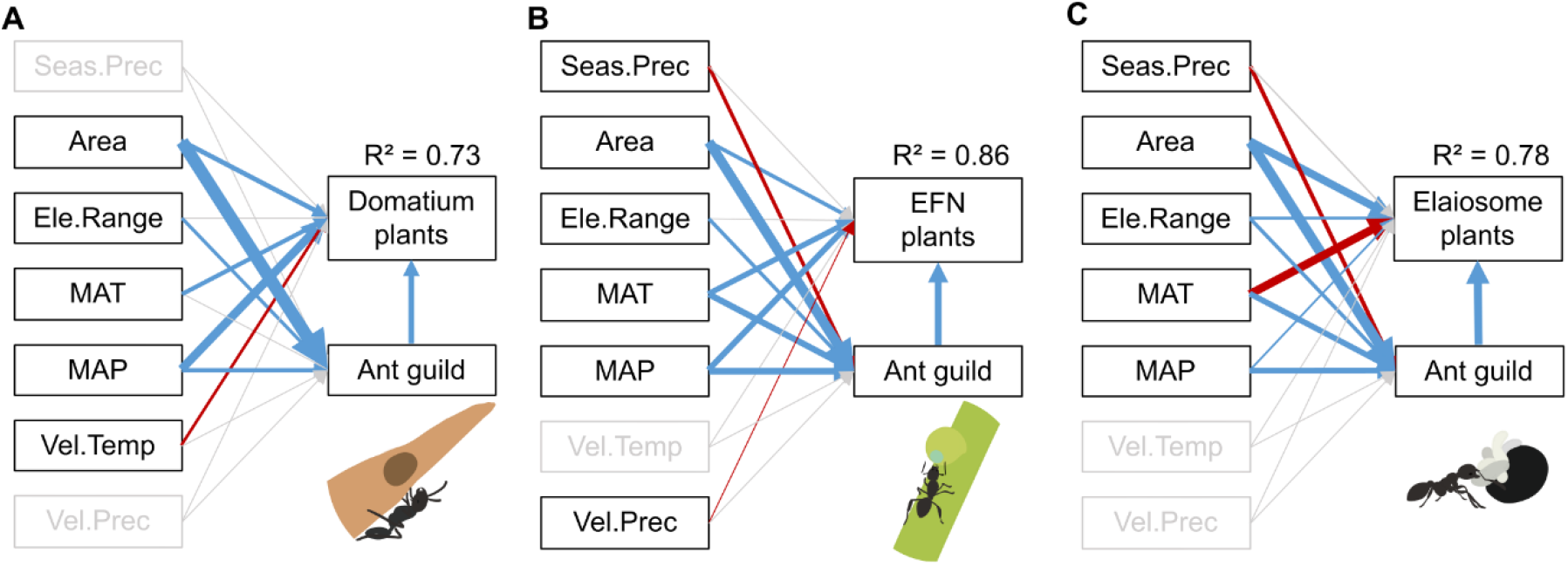
Structural equation models (SEMs) showing the effects of abiotic (climatic and geographic variables) and biotic (species richness of ant guilds) variables on richness patterns of (*A*) domatium-bearing, (*B*) extrafloral nectary (EFN)-bearing and (*C*) elaiosome-bearing plants. Boxes represent variables and arrows represent the direction of variable effects. Arrow size is proportional to the absolute value of path coefficients, blue and red arrows represent significant (p < 0.05) positive and negative path coefficients, respectively. Gray boxes and arrows represent non-significant (p > 0.05) variables. R^2^ is shown beside plant species richness. Ant guild, species richness of respective ant guild; Area, region area; Ele.Range, elevation range; MAT, mean annual temperature; MAP, mean annual precipitation; Seas.Prec, precipitation seasonality; Vel.Temp and Vel.Prec, climate-change velocity in temperature and precipitation from Last Glacial Maximum ∼21,000 y BP to present.

**Figure 3.**
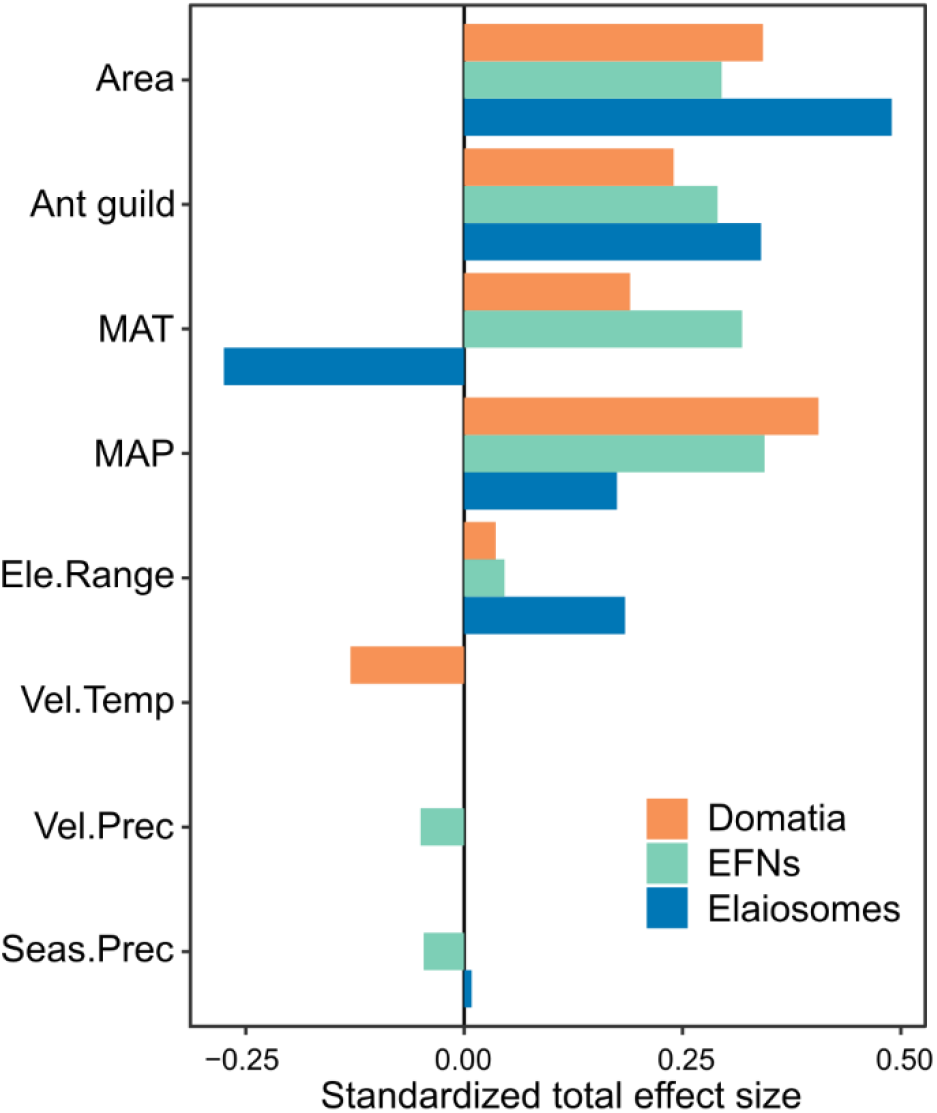
Standardized total effect sizes of abiotic and biotic predictors on species richness of domatium-, extrafloral nectary (EFN)- and elaiosome-bearing plants, calculated as the sum of direct and indirect path coefficients of structural equation modeling. Orange, green, and blue bars represent effects on domatium, EFN and elaiosome-bearing plants, respectively. Variables follow as Fig. 2.

Abiotic predictors alone account for a large proportion of variation in plant richness patterns (assessed as pseudo R^2^), yet ant richness remains a significant predictor of plant richness after controlling for abiotic predictors (all p-values < 0.001) (*SI Appendix*, Table S2). The strength of ant-plant relationships varies among mutualism types (standardized path coefficients for richness of ant guilds = 0.24, 0.29, 0.34, respectively) (Fig. 2). Models including respective ant guild richness generally have a better model fit (i.e., lower Akaike information criterion, higher R^2^) than abiotic SEMs (only including abiotic factors) for each mutualism (R^2^ values increase from 0.68 to 0.73 for domatia model, from 0.80 to 0.86 for EFNs, from 0.69 to 0.78 for elaiosomes) (Fig. 2 and *SI Appendix*, Table S2). Geographic patterns of proportions of ant guilds (relative to all ants) show similar latitudinal gradients to respective interacting plants, except the EFN-related ant guild (*SI Appendix*, Fig. S3).

As zero inflation occurred in the model of domatium-bearing plants (observed versus expected zeros from simulations, p-value = 0.008), we applied two separate analyses including one binomial model (modeling absence-presence of domatia) and one count model (only including non-zero richness of domatium-bearing plants) (29). The count model was used in further analyses to obtain comparable results. Results are largely consistent between binomial and count models of domatium-bearing plants and show major effects of ant guild richness and contemporary climate (*SI Appendix*, Table S2). We performed sensitivity analyses to evaluate whether the relationship between ants and plants are robust to ant guild assignment. When we applied species richness of total ants, ant guilds without hyperdiverse genera or trait-based ant guilds as biotic drivers, the overall associations of ants and interacting plants were consistent with respect to ant guild assignment (*SI Appendix*, Fig. S4).

## Discussion

Our study reveals distinct geographic patterns of plants with domatia, extrafloral nectaries (EFNs) and elaiosomes, which are generally consistent with empirical knowledge from previous studies at smaller spatial scales. Structural equation models show that these patterns are jointly driven by contemporary climate and species richness of respective ant guilds, yet no factor alone is sufficient in explaining the spatial variation of ant-associated plants. Although the effects of climate vary across different ant-plant mutualisms, our results highlight the consistent importance of both abiotic and biotic factors in shaping the diversity of ant-associated plants.

### Geography of ant-associated plants

Plants bearing domatia and EFNs show opposing geographic patterns to those bearing elaiosomes, with species richness of domatium- and EFN-bearing plants decreasing away from the equator while elaiosome-bearing plants are most diverse at mid-latitudes. The latitudinal gradients of plants with domatia and EFNs coincide with that of herbivory intensity (30), possibly because indirect defenses by domatia and EFNs are important components of plant-herbivore interactions (14, 31). On the contrary, as small seeds are portable and preferred by ants, the high prevalence of ant-dispersed plants at mid-latitudes may contribute to the observed pattern of decreasing seed mass at the edge of the tropics (32). Indeed, presence of domatia, EFNs and elaiosomes can correlate with other traits that may collectively mediate species responses to the environment (2).

Aside from the well-known diversity hotspots of elaiosome-bearing plants in South Africa and Australia (10, 25), the Mediterranean basin is expected to have more examples of ant-dispersed species than currently known, as it shares a similar climate to the former regions. We find surprisingly high diversity of ant-dispersed plants in the Mediterranean basin, as well as northern hemisphere temperate regions (e.g., Sichuan, China), where limited attention has been given to myrmecochory. In temperate forests, such as Central European and eastern North American forests, many ant-dispersed plants bloom in spring to differentiate from seed dispersal by birds (10). Interestingly, elaiosome-bearing plants are scarce in the Neotropics, although ants are highly diverse and reported in different phases of seed dispersal (e.g., ants may disperse one quarter woody species in the Caatinga but only 12.8% of them bearing elaiosomes on seeds) (33). While it is likely that sampling effort influenced this pattern, the low proportion of ant dispersers in our results confirm the scarcity of plants with elaiosomes in the Neotropics.

### Global drivers of ant-associated plant distributions

Temperature and precipitation positively affect species richness patterns of domatia and EFNs. Particularly, the tropics hold more ant-defended plant species than temperate or boreal regions, a phenomenon previously observed across regions (10). Temperature and precipitation may directly influence the ability of plants to produce sugary nectar, which attracts ants to establish mutualistic interactions (34). Likewise, ants are thermally sensitive and activity of ants against herbivores can be temperature-dependent (24, 35), leading to indirect effects via ants on the diversity of plants with domatia and EFNs. These may partly explain the low presence of ant-defended plants in temperate regions.

By contrast, temperature has a strong negative effect on elaiosome-bearing plants, while the effect of precipitation is significant but mild. These findings align with the high prevalence of elaiosomes in regions with harsh climates. Elaiosomes themselves may play an important role in increasing seed resistance to dry and freezing environments (36). Furthermore, ants disperse and store seeds in their nests, which may contribute to persistent soil seed banks (37). Seeds are therefore protected from fires, desiccation, and predation of rodents in suitable habitats for germination (38). Consequently, these unique benefits of myrmecochory are important for plants to persist in harsh environments that are relatively dry, with a high temperature range, and low soil nutrients (9, 38). In contrast, myrmecochory and its advantages might be unnecessary for plants in the Neotropics, which is in line with our results of low diversity of elaiosomes-bearing plants in this realm.

Additionally, region area has a positive effect on ant-associated plant diversity with the effect being not only direct, but even more indirectly mediated through ant diversity. Surprisingly, elevation range was a poor predictor of domatium- and EFN-bearing plant richness, possibly because temperature exerts a stronger influence on ant-defended plants and ant distributions. Likewise, precipitation seasonality has only minor negative effects on ant-associated plant richness indirectly via ant guild richness, indicating a potential risk of desiccation for ants in regions with high rainfall seasonality (35). Past climate change is generally influential on plants and ants (39, 40), however, we find a negligible importance on ant-associated plants. This is likely due to the divergent histories of biogeographic regions in our study, highlighting the need for a better understanding of biotic interactions under past climate change.

Besides abiotic conditions, species richness of ant guilds related to ant-plant mutualisms is an important factor associated with diversity patterns of domatium-, EFN- and elaiosome-bearing plants. The “ant-limitation hypothesis” (17) predicts that spatial variation of interacting plants may be affected by ant community attributes, such as ant abundance (41), activity (18) and species richness (15). This hypothesis has been proposed for processes acting at local communities. However, our findings lend support for this hypothesis at broader spatial scales, highlighting the importance of considering biotic interactions in species distributions and diversity across spatial scales (42). We also note some mismatch between the distributions of plants and related ant guilds. One mismatch is that domatium-related ants appear outside the tropics while domatia seldom occur in temperate regions. Ant-associated traits and mutualistic interactions can alter the response of plants to the environment by providing suits of services. Interacting ant richness does not affect ant-associated plant richness alone, but in combination with climate. Therefore, domatium-related ants are necessary but not sufficient to link to domatium-bearing plants, only when regions are warm and moist.

In addition, incomplete knowledge on ant ecology at the species level and especially interactions with plants, also hamper the precise estimation of geography and drivers of ant-plant mutualisms. Likewise, species richness of all three ant guilds is highly correlated to total ant richness (Pearson’s r > 0.9), as ant guilds are divisions of the same taxonomic group. Nonetheless, the proportion of the three ant guilds to total ants exhibits similar patterns to ant-associated plants, and our results are robust to the assignment of ant guilds as shown in our sensitivity analyses. Further refinements of ant distributions and ant-plant networks hold a great potential to promote our understanding of ant-plant mutualisms.

In summary, we reveal the direct and indirect drivers of global patterns of species richness of plants bearing domatia, EFNs and elaiosomes. Climate and associated ant diversity are essential to the geography of ant-associated plants, with the effects being mediated by features of mutualistic interactions. The consideration of mutualistic interactions together with abiotic factors is critical in deciphering the factors governing the global distribution and biogeography of plants.

## Materials and Methods

### Plant checklists and distributions

Information about flowering plant species involved in ant-plant mutualisms was sourced from comprehensive reviews on domatium-bearing plants (13), extrafloral nectary (EFN)-bearing plants (26) and elaiosome-bearing plants (11), which provide information about the presence of domatia and EFNs on plants at the species level and elaiosomes at the genus level. Before extracting checklists for further analyses, we omitted duplicates and taxonomically uncertain records, and standardized all plant names according to The Plant List v. 1.1 (43). This step led to 675 valid domatium-bearing species and 3275 valid EFN-bearing species. The elaiosome-bearing plant list contained 239 genera and for most genera all constituent species were dispersed by ants (11). We considered all species from myrmecochorous genera as elaiosome-bearing plants, including approximately 11,040 species.

Native occurrences of angiosperm species were retrieved from the Global Inventory of Floras and Traits database (GIFT 2.1) (28), which contains regional checklists of plants including 352,232 taxonomically standardized plant species across 3,088 regions worldwide. We only used non-overlapping regions with available plant occurrences. Most regions are political units (e.g., countries or administrative units) and geographic regions (e.g., islands). We excluded oceanic islands from this analysis due to their peculiar diversity and assembly patterns (44). Regional occurrences were then linked to the lists of domatium-, EFN- and elaiosome-bearing plants to obtain distribution information of plants interacting with ants. In total, our dataset contains 406 regions worldwide, including 373 mainland regions and 33 continental islands (with areas larger than 1000 km^2^).

### Ant guilds and distributions

Ants that may be involved in ant-plant mutualisms were determined from an exhaustive literature search and expert opinions (*SI Datasets* S1). We assembled three genus-level ant guilds, which are groups of ants that may interact with domatia, EFNs, and elaiosomes, respectively. Some genera were removed because species within these genera may play a dual role in interactions (sometimes as exploiters rather than partners). For instance, granivorous ants such as from the *Messor* genus, harvest seeds for food and only disperse a small number of them (45). Finally, we retrieved native ant species occurrences from the Global Ant Biodiversity Informatics database (GABI) (27), which includes more than 1.9 million distributional records of more than 15,700 ant species and subspecies from comprehensive publications, digitized museum collections and specimen databases. Ant occurrences were then linked with ant guilds to obtain species richness of plant-associated ants as biotic factors in our models.

### Abiotic variables

Past and present environmental factors are strong constraints acting on broad-scale diversity patterns of many taxa (46, 47). Water- and energy-related variables, as well as their fluctuations, may exert direct effects on the distribution and diversity of plants as well as indirect effects via ants (34, 39). We derived contemporary climate factors from the Climatologies at high resolution for the earth’s land surface areas dataset (CHELSA V1.2) (48), including mean annual temperature (°C, hereafter temperature), mean annual precipitation (mm, hereafter precipitation) and precipitation seasonality (standard deviation of the monthly precipitation). Elevation range (m, the maximum elevation minus the minimum elevation of a region), as a proxy for environmental heterogeneity (49), was derived from the Global multi-resolution terrain elevation data at a resolution of 30 arc sec (50). To control for differences in area size and subsequent species-area relationship, we further included area (km²) of each region as a covariate (46). Past climate stability has been documented to shape modern biodiversity patterns of plants and other taxa (47). Therefore, we also included past climate change velocity in temperature and precipitation, which were obtained as velocity of climate change in temperature and precipitation from the Last Glacial Maximum (LGM) 21,000 y BP to the present (47, 51, 52). Altitude, temperature seasonality, isothermality and aridity were initially considered yet were not included, as they showed significant correlations (Pearson’s r > 0.7) with other variables and weaker relationships with the richness of ants and plants. We derived all abiotic variables for each geographic region from the GIFT database (28).

### Modeling

Before data analysis, all abiotic variables, plant, and ant richness were log-transformed to improve the normality of model residuals, and z-transformed to have zero mean and unit variance. All statistical analyses were conducted in R version 4.1.1 (53).

To quantify the effects of environmental and biotic drivers on species richness of ant-associated plants, we fitted regression models of plant species richness using area, elevation range, present-day climate (temperature, precipitation, and precipitation seasonality), paleoclimate (past climate change velocity in temperature and precipitation) and species richness of ant guilds as predictors. As spatial autocorrelation was detected in model residuals (e.g., EFN-bearing plant: Moran’s I = 0.52, p-value < 0.001), we used simultaneous autoregressive (SAR) models of the spatial error type. Spatial models require a spatial weight matrix, which is based on the definition of neighborhood structure and spatial weights for each region. Because most of our regions are political units and inconsistent in geometry and size, we used the sphere of influence to identify the neighbors for each region, rather than using an adjacency or distance-band approach (54). The sphere of influence is defined as a circle around the centroid of a focal region within a radius equal to the distance to the centroid of the nearest region. Two regions were considered neighbors when the sphere of influence of these two regions overlapped. We then used row standardized coding to weigh the neighbors of each region. Spatial models performed well in dealing with our spatially autocorrelated data (e.g., EFN-bearing plant: Moran’s I = −0.08, p-value = 0.98). SAR models were implemented using the ‘spatialreg’ R package v. 1.1-5 (55).

To identify the direct and indirect effects of abiotic and biotic factors on plant richness, we fitted SAR models in a piecewise structural equation modeling (SEM) framework, which is appropriate to test multivariate causal hypotheses (56). Piecewise SEM is based on directed acyclic graphs and can incorporate many model structures (e.g., spatial correlation structure) (56). We first constructed a priori theoretical SEMs that included all hypothesized pathways among plants, ants, and abiotic factors. We examined the relationships in two SEMs: 1) abiotic SEMs, where plants and ants were not linked and only directly affected by abiotic variables; 2) full SEMs, where links were additionally specified from ants to plants. We fitted separate SEMs for domatia, EFNs and elaiosomes as the ant-plant mutualism. Goodness-of-fit was evaluated based on Shipley’s test of directed separation by comparing Fisher’s C statistic to the χ^2^ distribution, where an insignificant p value (p-value > 0.05) means no missing path exists (56). We optimized models in a stepwise manner, starting from removing the path with the lowest standardized path coefficient. The final models were determined by Fisher’s C statistic and Akaike information criterion (AIC). Nagelkerke’s pseudo R^2^ was calculated to evaluate model fit. SEMs were implemented using the ‘piecewiseSEM’ R package v. 2.0.2 (56).

To evaluate whether the relationships between plants and ant guilds are robust in our analysis, we performed sensitivity analyses by 1) using species richness of all ants as a biotic factor; 2) removing hyperdiverse genera (containing over 500 species per genus) in ant guilds; and 3) adapting another assignment of ant guilds based on ant ecological traits. Within hyperdiverse genera (i.e., *Camponotus, Pheidole, Polyrhachis, Crematogaster* and *Tetramorium*), some species were observed to protect plants and disperse seeds while many were not. Considering these genera may overestimate interacting ant species richness and bias the model results, we removed hyperdiverse genera (accounting for 29% of total ant species) and reran models. For ant guild assignment by ecological traits, we chose four genus-level ecological traits of ants as indicators of ant-plant interactions (57, 58): diet (herbivore, omnivore, or predator), nesting location (subterranean, or arboreal), foraging location (subterranean, or arboreal), and stratum (arboreal, epigaeic, or hypogaeic). These three methods of classifying ants as biotic factors were only applied in sensitivity analyses, which show that our results are robust to the method of ant guild assignment. For more details on ant guild assignment and results of sensitivity analyses see the SI Appendix.

## Author Contributions

Y.L., A.T., H.K. designed research; Y.L. compiled data with contributions from B.G., E.E. (for ant dataset) and P.W., A.N., I., H.K. (for plant dataset); Y.L. analyzed data with assistance from A.T., P.W., B.G., H.K.; Y.L. wrote the paper with assistance from all authors.

## Acknowledgments

Y.L. acknowledges funding from the China Scholarship Council (CSC, grant number 201808330422), A.T. acknowledges support from the German Research Foundation (Deutsche Forschungsgemeinschaft, DFG project number 447332176), H.K. acknowledges funding from the German Research Foundation (Research Unit FOR 2716 DynaCom).

