## Supplementary text, Figures S1 to S4, Tables S1 to S2 for "Climate and ant richness explain the global distribution of ant-plant mutualisms"

### Supplementary Information Text

**Ant guilds based on ecological traits.** To represent potentially selective forces on the evolution of ant-associated traits in plants and interactions with plants (1-3), we chose four genus-level ecological traits of ants from previous literature (4, 5), as indicators of ant-plant interactions: diet (herbivore, omnivore, or predator), nesting location (subterranean, or arboreal), foraging location (subterranean, or arboreal), and stratum (arboreal, epigaeic, or hypogaeic). Based on ant behaviors related to interactions, we then classified three ant guilds corresponding to three interaction types: 1) ant guild associated with domatia is characterized by arboreal nesting and foraging ants, and occupy arboreal habitats (3); 2) guild associated with EFNs is typically represented by predacious ants that forage arboreally (3, 6); 3) guild interacting with elaiosomes is ant dispersers that are omnivores, forage and nest on the ground, and occupy epigaeic habitats (2). Some genera were considered as polymorphic or ambiguous due to the evidence of alternative trait states or the absence of documented records, making up 176 of 338 ant genera and about 58% of all ant species. For example, species of *Pheidole* have highly diverse nesting preferences and were assigned nesting polymorphism. Therefore, we opted for a conservative approach and thus categorized polymorphic and ambiguous traits to each ant guild. For example, as species of *Pheidole* are omnivorous and polymorphic in nesting and foraging locations, we categorized *Pheidole* to both guilds associated with EFNs and domatia.

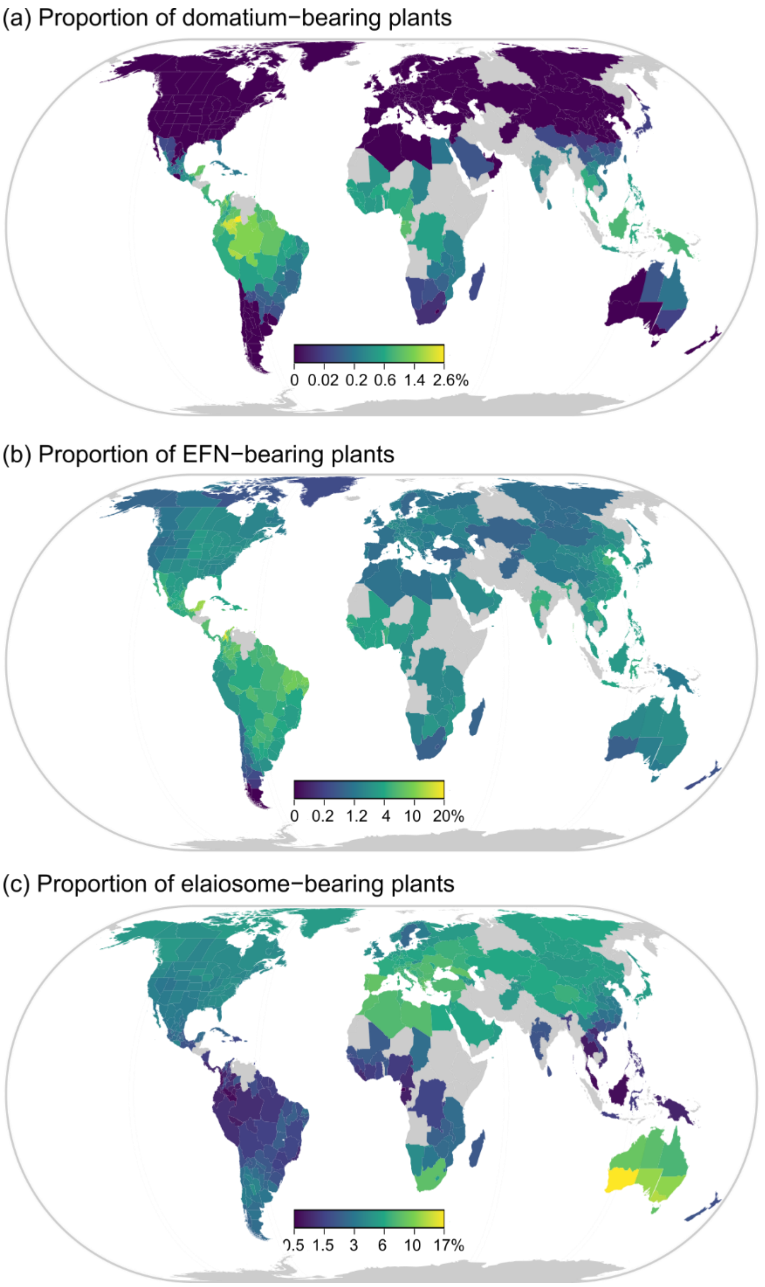


**Fig. S1.** Global patterns of the proportional representation of plants with (a) domatia, (b) EFNs and (c) elaiosomes. Proportion is ant-associated plant species richness to angiosperms per region. Gray color indicates regions without data.

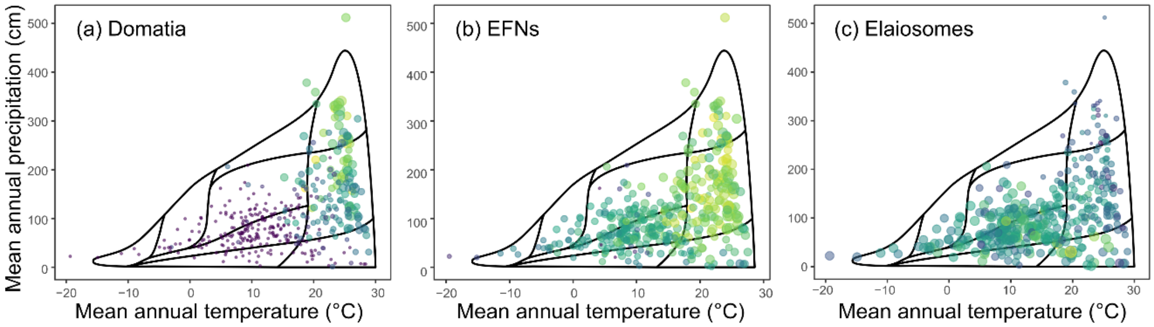


**Fig. S2.** Distributions of plants with (a) domatia, (b) EFNs and (c) elaiosomes in environmental space, overlaid by Whittaker's biome. The color of the circles follows the species richness schemes in Figure 1, respectively. The size of the circles is corresponding to the proportion of ant-associated plant species richness to angiosperms per region.

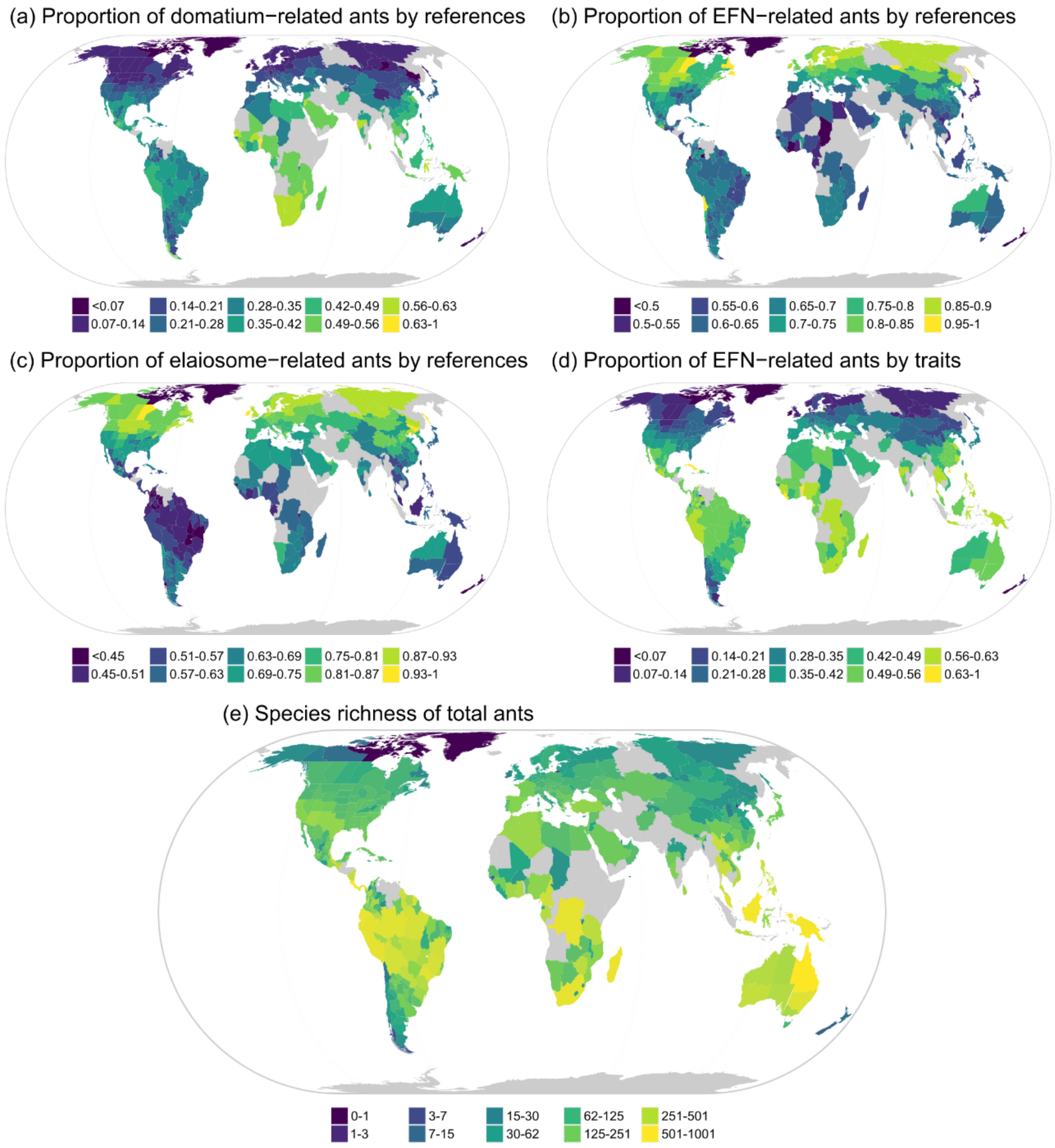


**Fig. S3.** Global patterns of proportional representation for ant guilds interacting with (a) domatia, (b) EFNs, (c) elaiosomes based on references, (d) elaiosomes based on traits, and (e) species richness for total ants. Proportion is ant-associated plant species richness to angiosperms per region. Gray color indicates regions without data. For ant diversity maps, also see antmaps.org.

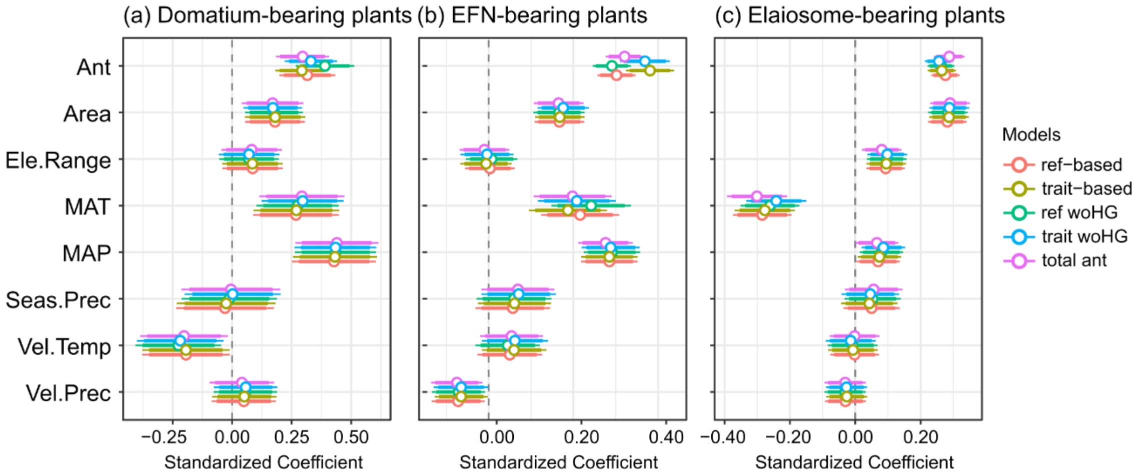


**Fig. S4.** Sensitivity analyses for plants bearing (a) domatia, (b) extrafloral nectaries (EFNs), and (c) elaiosomes. Ref-based, refers to reference-based ant guild assignments, which are applied in our main results; trait-based, refers to trait-based ant guild assignments; ref woHG, refers to reference-based ant guild assignments removing hyperdiverse genera; trait woHG, refers to trait-based ant guild assignments removing hyperdiverse genera; total ant, refers to total ant species richness as an explanatory variable of biotic interactions. Ant, species richness of respective ant guild; Area, region area; Ele.Range, elevation range; MAT, mean annual temperature; MAP, mean annual precipitation; Seas.Prec, precipitation seasonality; Vel.Temp and Vel.Prec, climate-change velocity in temperature and precipitation from Last Glacial Maximum ~21,000 y BP to present.

**Table S1.** Species richness and proportional representation of plants bearing domatia, EFNs and elaiosomes in top 5 regions.

| | Regions in richness | Species richness | Regions in proportion | Proportion |
|---|---|---|---|---|
| Domatia | Peru | 118 | Guainía, Colombia | 0.025 |
| | Amazonas, Brazil | 107 | Caquetá, Colombia | 0.020 |
| | Ecuador | 99 | Guaviare, Colombia | 0.020 |
| | Thailand | 93 | Vaupés, Colombia | 0.019 |
| | New Guinea | 91 | Amazonas, Colombia | 0.017 |
| EFNs | Bolivia | 513 | Sucre, Colombia | 0.193 |
| | Peru | 509 | Atlántico, Colombia | 0.168 |
| | Costa Rica | 459 | Bolívar, Colombia | 0.131 |
| | Panama | 457 | Córdoba, Colombia | 0.127 |
| | Chiapas, Mexico | 456 | Quintana Roo, México | 0.119 |
| Elaiosomes | South Africa | 1528 | South Western Australia | 0.167 |
| | South Western Australia | 1243 | Victoria, Australia | 0.130 |
| | Sichuan, China | 684 | South Australia | 0.117 |
| | New South Wales, Australia | 641 | New South Wales, Australia | 0.114 |
| | Asiatic Turkey | 640 | North Western Australia | 0.094 |

**Table S2.** Model results of domatia (binomial & count), EFNs and elaiosomes. AIC and R-squared of models without predictor of ant are indicated in brackets. Domatia of the binomial model was applied by a generalized linear model with binomial distribution, and the rest three models were applied by spatial autoregressive models with error type. Ant, species richness of related ant guild; Area, region area; Ele.Range, elevation range; MAT, mean annual temperature; MAP, mean annual precipitation; Seas.Prec, precipitation seasonality; Vel.Temp and Vel.Prec, Last Glacial Maximum climate-change velocity in temperature and precipitation.

| Models | Variables | Estimate | Std.Error | P value | AIC | R-squared |
|---|---|---|---|---|---|---|
| Domatia (binomial) | Ant | 2.13929 | 0.616701 | *** | 106 (120) | - |
| | Area | 0.773221 | 0.373498 | * | | |
| | Ele.Range | -0.3271 | 0.365796 | | | |
| | MAT | 1.448331 | 0.43356 | *** | | |
| | MAP | 1.669134 | 0.433067 | *** | | |
| | Seas.Prec. | 1.570406 | 0.539536 | ** | | |
| | Vel.Temp | -0.94733 | 0.535691 | | | |
| | Vel.Prec | 0.374044 | 0.446713 | | | |
| | Autocov. | 4.594882 | 1.051433 | *** | | |
| Domatia (count) | Ant | 0.316078 | 0.059899 | *** | 306 (329) | 0.73 (0.68) |
| | Area | 0.179597 | 0.06447 | ** | | |
| | Ele.Range | 0.086107 | 0.065002 | | | |
| | MAT | 0.268111 | 0.091533 | ** | | |
| | MAP | 0.427529 | 0.089501 | *** | | |
| | Seas.Prec. | -0.0296 | 0.105386 | | | |
| | Vel.Temp | -0.19423 | 0.093582 | * | | |
| | Vel.Prec | 0.04901 | 0.068857 | | | |
| EFNs | Ant | 0.301959 | 0.022953 | *** | 390 (532) | 0.86 (0.80) |
| | Area | 0.166697 | 0.03045 | *** | | |
| | Ele.Range | 0.003013 | 0.030291 | | | |
| | MAT | 0.216374 | 0.047441 | *** | | |
| | MAP | 0.284592 | 0.033828 | *** | | |
| | Seas.Prec. | 0.056962 | 0.044549 | | | |
| | Vel.Temp | 0.050078 | 0.038587 | | | |
| | Vel.Prec | -0.07203 | 0.031609 | * | | |
| Elaiosomes | Ant | 0.275684 | 0.022047 | *** | 386 (516) | 0.78 (0.69) |
| | Area | 0.282217 | 0.030473 | *** | | |
| | Ele.Range | 0.092524 | 0.030436 | ** | | |
| | MAT | -0.28456 | 0.046248 | *** | | |
| | MAP | 0.06967 | 0.033745 | * | | |
| | Seas.Prec. | 0.049868 | 0.044021 | | | |
| | Vel.Temp | -0.00249 | 0.038456 | | | |
| | Vel.Prec | -0.02956 | 0.031675 | | | |

**Dataset S1 (separate file).** Database of ant guild assignments for 338 ant genera, based on an exhaustive literature search and expert opinions.

**Dataset S2 (separate file).** Database of diversity of domatium-, extrafloral nectary- and elaiosome-bearing plants and related ant guilds on 406 regions, in coupled with abiotic predictors.
